# Automatic identification of bird females using egg phenotype

**DOI:** 10.1101/2020.11.25.398131

**Authors:** Michal Šulc, Anna E. Hughes, Jolyon Troscianko, Gabriela Štětková, Petr Procházka, Milica Požgayová, Lubomír Piálek, Radka Piálková, Vojtěch Brlík, Marcel Honza

## Abstract

1. Identification of individuals greatly contributes to understanding animal ecology and evolution, and in many cases can only be achieved using expensive and invasive techniques. Advances in computing technology offer alternative cost-effective techniques which are less invasive and can discriminate between individuals based on visual and/or acoustic cues. Here, we employ human assessment and an automatic analytical approach to predict the identity of common cuckoo (*Cuculus canorus*) females based on the appearance of their eggs. The cuckoo’s secretive brood parasitic strategy makes studying its life history very challenging. Eggs were analysed using calibrated digital photography for quantifying spotting pattern, size and shape, and spectrometry for measuring colour. Cuckoo females were identified from genetic sampling of their nestlings, allowing us to determine the accuracy of human and automatic female assignment. Finally, we used a novel ‘same-different’ approach that uses both genetic and phenotypic information to assign eggs that were not genetically analysed.
2. Our results supported the ‘constant egg-type hypothesis’, showing that individual cuckoo females lay eggs with a relatively constant appearance and that eggs laid by different females differ more than eggs laid by the same female. The accuracy of unsupervised hierarchical clustering was comparable to assessments of experienced human observers. Supervised random forest analysis showed better results, with higher cluster accuracy. Same-different analysis was able to assign 22 of 87 unidentified cuckoo eggs to seven already known females.
3. Our study showed that egg appearance on its own is not sufficient for identification of individual cuckoo females. We therefore advocate genetic analysis to be used for this purpose. However, supervised analytical methods reliably assigned a relatively high number of eggs without genetic data to their mothers which can be used in conjunction with genetic testing as a cost-effective method for increasing sample sizes for eggs where genetic samples could not be obtained.

## Introduction

Identification of individuals is important in zoological research, particularly when investigating variation among or within individuals in a population. Traditionally, capture-mark-recapture techniques have been used to monitor individuals during their lifetime (Lindberg, 2012; Jung, Boonstra, & Krebs, 2020). This method has been improved by employing more sophisticated methods such as attaching GPS (global positioning system) and radio transmitters or RFID (radio frequency identification) tags (Krause et al., 2013) that allow researchers to investigate the spatial activity of animals in more detail. However, these methods still require capturing and tagging that is usually time-consuming, expensive (depending on the method used), and may reduce animal welfare (Weinstein, 2018). Therefore, cost-effective indirect approaches have been developed to identify and monitor individuals within the same species.

These indirect approaches rely on the fact that individuals differ from each other visually or acoustically and this variation may be used for their identification. Indeed, it has been shown that e.g. face (Deb et al., 2018; Hansen et al., 2018; Hou et al., 2020) and body pattern data (Hiby et al., 2009; Bolger, Morrison, Vance, Lee, & Farid, 2012; Crall, Stewart, Berger-Wolf, Rubenstein, & Sundaresan, 2013; Ferreira et al., 2020) captured from photographs may allow discrimination of individuals. Similarly, sounds produced by animals (especially bird song) also seem to serve as a good individual fingerprint (Blumstein et al., 2011; Petrusková, Pišvejcová, Kinštová, Brinke, & Petrusek, 2016; Ptacek, Machlica, Linhart, Jaska, & Muller, 2016). Recently, applying modern computer technology and artificial intelligence techniques (such as convolutional neural networks) that automate the analysis of various types of data from different sources such as pictures or audio recordings has made these methods reliable and applicable for various animal taxa (Hansen et al., 2018; Christin et al., 2019; Ferreira et al., 2020; Hou et al., 2020).

However, for many species, identification of all individuals in a population is still not straightforward e.g. due to their secretive behaviour or due to the fact that it is difficult to catch them. Here, we focus on one group of animals that are especially challenging to study – avian brood parasites. There are more than a hundred obligate brood parasites that never build their own nests and instead lay their eggs into nests of other species (Davies, 2010). Even more species belong to conspecific brood parasites that only occasionally lay eggs into nests of the unrelated conspecifics (Bruce E. Lyon & Eadie, 2008). Brood parasites and their hosts have been the focus of considerable research into co-evolutionary arms races (Soler, 2017). But since brood parasites only lay eggs and then usually do not return to host nests (but see Šulc et al. 2020), and because egg laying is (especially in obligate brood parasites) very quick (McMaster, Neudorf, Sealy, & Pitcher, 2004; Gloag, Fiorini, Reboreda, & Kacelnik, 2013; Jelínek, Šulc, Štetková, & Honza, 2020), direct observation of parasitism in nature is difficult and identification of parasitic females is problematic. As a consequence, many important aspects of females’ life history strategy are still poorly understood; in obligate brood parasites, we e.g. still know little about spatio-temporal distribution of their egg laying, consistency in host selection and the total number of eggs they lay during a breeding season. Conspecific brood parasitism is even less understood and we do not even know why some females adopt this strategy (Bruce E. Lyon & Eadie, 2008).

The idea of identifying bird females according to the appearance of the eggs they laid depends on the presumption that within-clutch variation in egg appearance is lower than between-clutch variation which has been confirmed for several species including brood parasites (Øien, Moksnes, & Røskaft, 1995; McRae & Burke, 1996; Paillisson, Latraube, Marion, & Bretagnolle, 2008; Höltje, Mewes, Haase, & Ornés, 2016). This approach has therefore been applied (although with caveats) for the identification of parasitic eggs in some conspecific brood parasite species (Gibbons, 1986; Møller, 1987; Jackson, 1992; Petersen, 1992; Bruce E. Lyon, 1993; McRae & Burke, 1996; B. E Lyon, 2003). However, some studies that estimated accuracy of parasitic egg identification showed ambiguous results for some species (Ådahl, Lindström, Ruxton, Arnold, & Begg, 2004; Pöysä, Lindblom, Rutila, & Sorjonen, 2009; Eadie, Smith, Zadworny, Kühnlein, & Cheng, 2010; Lemons, Sedinger, & Randle, 2011; Petrželková, Pöysä, Klvaňa, Albrecht, & Hořák, 2017) and for others this method did not work at all (Brown & Sherman, 1989; Cariello, Lima, Schwabl, & Macedo, 2004; Grønstøl, Blomqvist, & Wagner, 2006; Griffith, Barr, Sheldon, Rowe, & Burke, 2009; Roy, Parker, & Gates, 2009). One of the reasons why many studies found low accuracy of identification might be that closely related females lay indistinctive eggs. Several studies showed that egg appearance, namely egg color (Wei, Bitgood, & Dentine, 1992; Collias, 1993; Morales et al., 2010), spotting pattern (Gosler, Barnett, & James Reynolds, 2000) and egg size (Christians, 2002) are highly heritable traits which might complicate such analyses especially in inbred populations. Another explanation might be that previous studies did not use the full potential of egg variability (e.g. none of the presented studies measured egg colour in the ultraviolet range of spectrum).

Identification of parasitic females using egg appearance has also been attempted in the common cuckoo (hereafter cuckoo), but was unsuccessful (Moksnes et al., 2008). However, the study assessed cuckoo eggs from a human perspective, with people sorting the eggs based on photographs. To date, there have been no attempts to use more objective quantification methods for egg classification in the cuckoo. These objective methods, such as spectrophotometry for measuring colours (including the ultraviolet part of the spectrum), and image analysis of photographs for quantifying spotting pattern, size and shape of eggs are now available, and may allow more accurate classification that can be carried out in an automated manner.

In this study, we employ a novel automatic analytical approach to analyse phenotypic features of cuckoo eggs such as dimensions (size, shape), colour and spotting pattern to predict maternal identity. If successful, this low-cost and minimally invasive female identification method would greatly facilitate studies into a range of key questions regarding this secretive brood parasitic species. We also performed human assessment based on sorting of photos with cuckoo eggs to compare the reliability of both methods with the true identity acquired from molecular analyses. Moreover, we believe this automatic technique might be also used in other brood parasitic systems or in species where females are difficult to catch (see e.g. Höltje et al. 2016). Finally, it has been suggested that similarly looking eggs laid by different cuckoo females may belong to closely related females, e.g. mother and daughter (Moksnes et al., 2008). Therefore, we will for the first time investigate the relationship between the genetic distance of individual cuckoo females and the phenotypic distance of their eggs.

## Material and Methods

### Study system and data collection

All data were collected in the fishpond area between Mutěnice (48°54′N, 17°02′E) and Hodonín (48°51′N, 17°07′E) in South Moravia, Czech Republic from May to July 2017. We systematically searched the littoral vegetation for the great reed warbler (*Acrocephalus arundinaceus*) and Eurasian reed warbler (*Acrocephalus scirpaceus*) nests. Most great reed warbler (hereafter GRW) nests were found during the building stage when mapping male territories and mating status (Bensch, 1996). The rest of the GRW and all Eurasian reed warbler (hereafter RW) nests were found in different stages of breeding by systematic searching. If possible, all GRW nests were checked every day from the nest building stage until clutch completion and approximately every third day during incubation. All RW nests were checked approximately every second day during laying stage and extensively during incubation. GRWs experienced 92 % (59 out of 64 nests) and RWs 20 % (91 out of 456 nests) cuckoo parasitism rate. Multiple parasitism was also common; 37 of 59 and two of 91 parasitized GRW and RW nests, respectively, were parasitized by more than one cuckoo egg.

When a cuckoo egg was found in a host nest, we immediately measured its colour and took a photo (see below) to avoid colour change during the incubation period (Hanley et al., 2016). In the cases of multiply parasitized nests, we removed the newly laid cuckoo egg(s), transferred them to an incubator (HEKA-Kongo; HEKA-Brutgeräte, Rietberg, Germany) and incubated them artificially to prevent sample losses caused by the cuckoo nestlings (Honza, Vošlajerová, & Moskát, 2007). The removed cuckoo eggs were either incubated until hatching and chicks placed into non-parasitized host nests (for other experiments) or we froze the eggs before hatching for the future genetic analysis (see *Genotyping and kinship analysis* section). We took a blood sample from all 10-days old cuckoo nestlings from their ulnar or medial tarsometatarsal vein (approx. 25 μl). Finally, we mist-netted 29 and 16 adult cuckoo males and females, respectively, and collected their blood samples from the ulnar vein (approx. 25 μl). All blood samples were stored in 96% ethanol until later genetic analyses.

Altogether we found 203 cuckoo eggs (121 and 82 in the GRW and RW nests, respectively). We photographed and measured the colour of 192 of them. Among these photographed cuckoo eggs, genetic samples were collected from 105 nestlings or embryos.

### Measurements of egg appearance

To obtain background colour we measured reflectance using JAZ Spectrometer (Ocean Optics, Dunedin, FL, U.S.A.) in the range 300–700 nm, as that is the wavelengths range birds can perceive (Cuthill 2006). We took nine measurements (each covering approximately 1 mm^2^) at three different parts of the egg (sharp pole, middle part and blunt pole). Since we focused on background colour, we avoided measuring dark spots (Šulc et al. 2016). For each egg, we used the measurement with the highest reflectance that best corresponded to the colour of the background (Šulc et al. 2019).

Spotting pattern was calculated from digital images taken by a Canon EOS 700D with prime Canon EF 40 mm lens. All photos were taken under standardized diffuse sunlight conditions (using a photography light tent), at the same angle and from the same distance and were referred to a grey standard (X-Rite Colour Checker Grey Scale Chart) with known reflectance. Image calibration, pattern analysis, analysis of shape and measurements of size were performed in ImageJ software (Schneider et al. 2012) using the Multispectral Image Calibration and Analysis (MICA) Toolbox (van den Berg et al. 2020). A scale bar was included in each photo, meaning that all images were equally rescaled to the scale of the smallest image (30 pixels/mm), and egg dimensions were obtained from the photos. For pattern investigation we applied a granularity analysis approach (Troscianko and Stevens 2015) that creates a bandpass ‘energy’ spectrum across a range of spatial frequencies, and then the pattern energy at each frequency band was measured as the standard deviation of the filtered image (for details see Šulc et al. 2019 and van den Berg et al. 2020). Since pattern energy does not discriminate the direction of the pattern (it cannot distinguish between dark spots on light background and light spots on dark background), we also calculated the ‘skew’ of the pattern, which quantifies the asymmetry of the pattern luminance distribution. A negative value of skew implies there are more spots than background colour, while a positive skew implies there is more background colour than spots. Skew was measured independently at each granularity band. All colour measurements and photos were taken by a single person (M.Š.) to ensure high consistency of the data.

### Genotyping and kinship analysis

DNA was isolated from the blood of adults and nestlings or tissues of embryos. (65 nestlings, 41 embryos, 29 adult males and 16 adult females). We estimated kinship relationships from nuclear SNPs and mitochondrial DNA haplotypes enabling us to exclude highly implausible maternal (or maternal-sibling) relationships in the inferred genealogy. As an additional criterion we also used the laying date of cuckoo eggs (data from daily checks) because it is known that a cuckoo female cannot lay her eggs more often that every second day (Chance, 1922; Payne, 1973; Wyllie, 1981).

To acquire the SNP dataset, we genotyped all samples with the ddRAD (double digest restrictionsite associated DNA) technique (Petersen, 1992) following the protocol of (Piálek et al., 2019). Two prepared libraries were sequenced on an Illumina HiSeq4000 system (2 lanes, 150 cycles P/E) in the EMBL Genomic Core Facility, Heidelberg, Germany. The obtained RAD-tags were processed in Stacks v2.4 (Catchen et al. 2011, Rochette et al. 2019) and mapped on the Cuculus canorus genome GCA000709325.1 (https://www.ncbi.nlm.nih.gov) with Bowtie2 assembler v2.2.4 (Langmead & Salzberg, 2012). Only loci with 95% or higher presence of individuals were scored and further filtered based on Hardy–Weinberg equilibrium, linkage disequilibrium and minimum minor allele frequency (0.4) in PLINK v1.9 (Purcell et al., 2007) which resulted in a dataset with 1620 markers. Kinship relationships were estimated using Colony (Jones & Wang, 2010) based on >1000 nuclear SNPs.

For the mitochondrial haplotype analysis, we sequenced a 411-bp portion of the left-hand hypervariable control region (Gibbs et al., 2000; Fossøy et al., 2011, 2012). Mitochondrial sequence data were assembled and manually checked in Geneious v10.2.6 (Kearse et al., 2012) and haplotypes were estimated based on a distance matrix with up to 1% tolerance (approx. 4 mutations) for genotyping errors.

Kinship analysis assigned the offspring (n = 105) to 30 clusters containing 1–10 eggs each (Fig. 1–5 in Supplementary material). Among these 30 clusters, nine corresponded to females that were caught and genotyped as described above. Thus, we were able to calculate genetic distances among these females.

**Figure 1.**
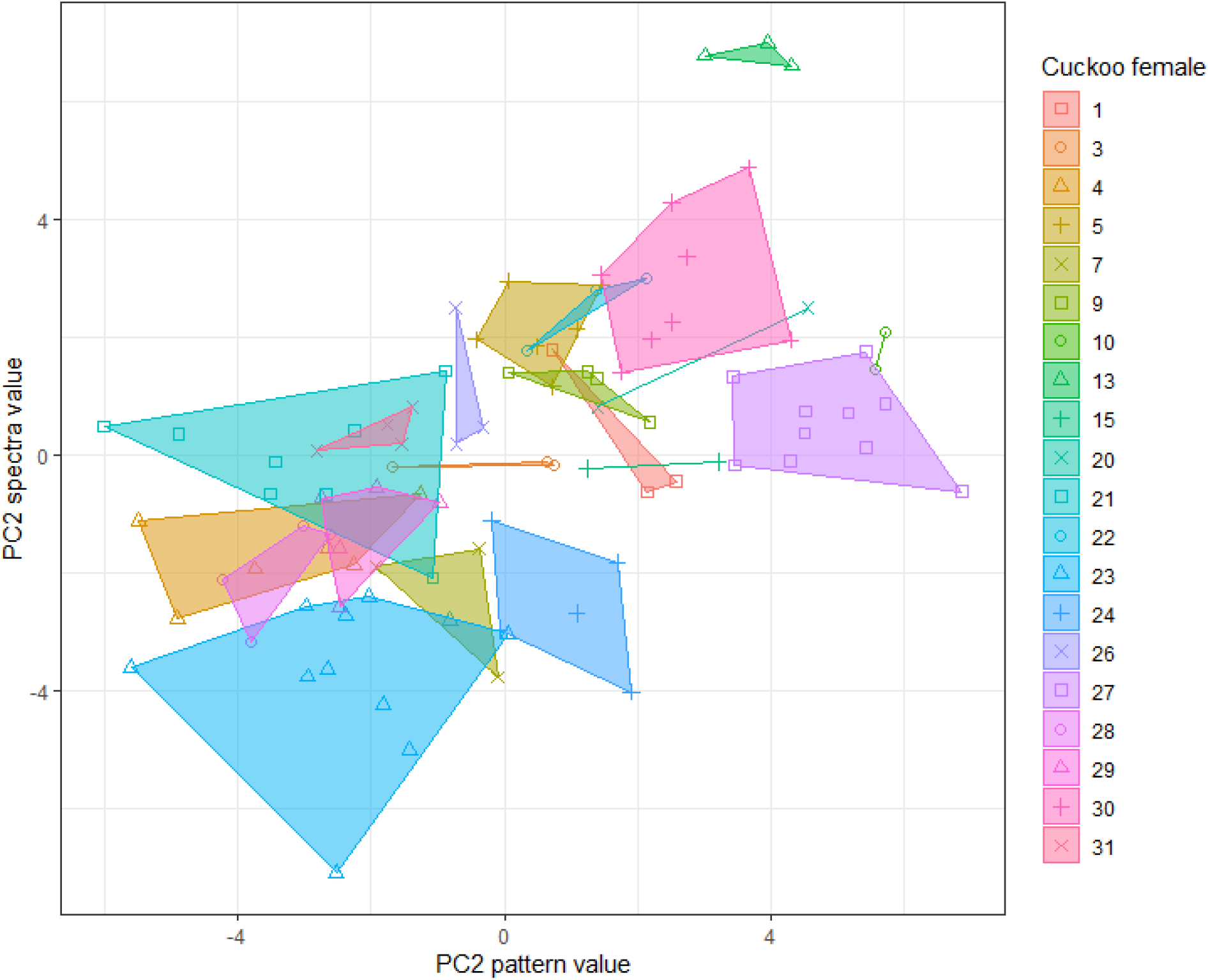
Values for individual eggs on the two most important PC variables (according to the random forest model), grouped by cuckoo female. PCA2 pattern variable indicates egg skew and PC2 spectra variable indicates blueness/greenness of eggs (for details, see Table 2).

For subsequent analysis dealing with egg phenotype (human and automatic assessment, see below), we removed females to which only one egg has been assigned (n = 10), meaning that we used a final dataset of 95 eggs laid by 20 females.

### Human assessment

We printed 95 photographs of cuckoo eggs that were standardized in their colour and size using the MICA Toolbox (van den Berg et al. 2020; Fig. 1–7 in supplementary material). We then asked twelve people to sort these photographs and create groups of pictures representing individual females according to similarity in egg appearance. Firstly, we asked them to sort these pictures into an unknown number of groups and, secondly, we asked them to sort these pictures into 20 groups corresponding to the real number of females identified by genetic assessment of identity. For the assessments, we asked 1) five people with no experience with egg appearance from wild animals, 2) three students of avian ecology that had experience with egg appearance from wild birds but had never seen cuckoo eggs and 3) four people (mostly co-authors of this manuscript) that had years of experience with cuckoo eggs. All participants received no other information about the eggs. Cluster similarity in egg classification was assessed using the adjusted Rand index, which provides a corrected-for-chance measure of the similarity between two data clusterings, implemented using the ‘cluster_similarity’ function from the R package clustereval (Ramey, 2012).

### Automatic assessment

We developed an automatic method based on the similarities/differences of cuckoo egg phenotypes. In the first step, we collected colour, pattern and dimension data from calibrated photographs and spectrophotometry data (for details, see Šulc et al., 2019) for all cuckoo eggs. Initially, we conducted Principal component analysis (PCA) on different aspects of the egg photographs, in order to avoid the use of correlated variables in the models.

#### Spectral data

a spectrophotometer was used to assess the background colour of cuckoo eggs (for details, see *Measurements of egg appearance*). PCA was carried out using binned, scaled spectral data created in the R package *pavo* (Maia, Gruson, Endler, & White, 2019), and two spectral PCA components were used in the final dataset (based on scree plot inspection). We also used two other spectral measures extracted from *pavo*: the mean brightness (B2; mean relative reflectance over the entire spectral range) (Hill & Mcgraw, 2006) and the position of the ultraviolet (UV) peak (defined as a wavelength within the range of 300–360nm where reflectance reached the highest point).

#### Shape data

the variables entered into the PCA were length, maximum width, volume, ellipse deviation and surface area (Troscianko, 2014). Three shape PCA components were selected for inclusion into the final dataset based on scree plot inspection.

#### Pattern data

the variables entered into the PCA were 12 pattern energies measured at a range of scales (from 1 to 0.0221 in steps of 1/square root of 2) across the whole egg (Troscianko & Stevens, 2015), and 12 pattern energy skew values measured at the same range of scales across the whole egg. We also included a measure of total pattern energy across the whole egg. Finally, we divided up each egg into three segments and measured the total pattern energy in each segment as well as the standard deviation between segments, to get a measure of how variable the patterning was across the egg. Three pattern PCA components were selected for inclusion into the final dataset based on scree plot inspection.

#### Luminance data

we analysed luminance from photographs, including both the spots and background areas of the eggs. We divided the egg up into three segments and took the average luminance and the standard deviation of luminance across each segment, as well as the standard deviation of luminance across all three segments. One luminance PCA component was selected for inclusion into the final dataset based on scree plot inspection.

In total, the final dataset contained 11 egg phenotypic traits that were used for clustering analysis.

### Within- and between-female variability in egg appearance

To create a metric of within-female variance, we calculated the standard deviation for each phenotypic trait within one female, and then took a mean value across all traits, giving an average variability value for each female.

To create a metric of between-female variance, we calculated the average value of each phenotypic trait for each female (i.e. created an “average” egg) and then calculated the standard deviation for each phenotypic trait across all females. We then averaged these standard deviations to create a measure of between-female variability across all traits. All trait values were scaled to ensure comparability across different traits.

To test whether within-female variance is lower than between-female variance, we conducted a one-sample t-test where the within-female variance metric (n=20) is compared with the test value (the between-female variance value).

### Unsupervised learning

Firstly, we carried out hierarchical clustering to attempt to cluster the eggs via visual similarity without any training or further information (e.g. number of females present). All variables were scaled for this analysis. To assess the accuracy of this method, we cut the tree by specifying the real number of groups (20) and assessed the cluster similarity between the predictions of the hierarchical model compared to the real data using the adjusted Rand index, as before.

### Supervised learning

#### Female clustering

We used a random forest model with a ‘leave-one-out’ cross-validation approach (Stone, 1974). For each egg in the dataset, the model was trained using a dataset consisting of all other eggs, and then tested using the focal egg. The model attempted to classify each egg to a given female, and the accuracy of the model was assessed using the classification accuracy value, and through cluster similarity values, as before (taking the average of 1000 runs, as random forest modelling is a stochastic process). We also fitted a random forest model to the full dataset to allow us to assess the importance of the different variables included in the model (using the mean decrease in accuracy).

#### Same/different analysis

We used an approach where a random forest model was trained to label pairs of eggs as ‘same’ or ‘different’. The training set used 3000 ‘same’ rows, where the two eggs were from the same female (but are not identical to each other) and 3000 ‘different’ rows, where the two eggs were from different females.

To test our models, we tested each egg in the labelled dataset on all eggs sequentially, including itself. We first tested whether the model recognised the identical eggs as being the same. We then tested whether each egg was only paired with other eggs from the same female i.e. whether the model could uniquely identify clusters of eggs that belonged together. The entire process (creating a training set, training the random forest model and testing the model) was repeated 1000 times.

For the unlabelled dataset, we calculated how many times in each of these 1000 runs the target egg was matched with a cluster of eggs that were from the same female. If the percentage was greater than 95%, we considered this egg as a candidate for being from this female. To corroborate this conclusion, we used non-phenotypic data e.g. laying dates, laying locality and host species.

### Phenotypic-genotypic similarity

We had genetic samples for 9 of the adult females, allowing us to create a genetic distance matrix. To compare the phenotypic-genotypic similarity between these females and their eggs, we created a trait distance matrix by taking means of the phenotypic parameters from their egg data, and then using Euclidean distance as the distance metric. We compared the genetic distance matrix with the trait distance matrix using a Mantel test, a statistical test of the correlation between two matrices, implemented in the vegan package in R using the Kendall method (as this is most appropriate for a small dataset). We also split the phenotype data into different components (spectral, pattern and shape) and calculated the phenotype-genotype similarities for each of these components separately, to test whether different aspects of the egg phenotype are differentially correlated with the female genotypes.

## Results

### Within- and between-female variability in egg appearance

Within-female variance was variable, with some females having very similar eggs (e.g. female 13 - within-female variance = 0.33, Fig. 2 in Supplementary material) and others having relatively high variance (e.g. female 29 - within-female variance = 1.31, Fig. 4 in Supplementary material). The between-female variance (comparing across females, using an “average egg” for each female) was 1.832 (mean of trait standard deviations, n = 11 traits; SD = 1.021). Overall, within-female variance (mean = 0.850, SD = 0.295) was smaller than between-female variance (one sample t test, t = 14.87, df = 19, p < 0.001). Variability in the egg appearance is also visible in Fig. 1 where the two most informative variables in the random forest analysis (PC2 for pattern and PC2 for spectral data), are plotted (see below and Table 2).

**Figure 2.**
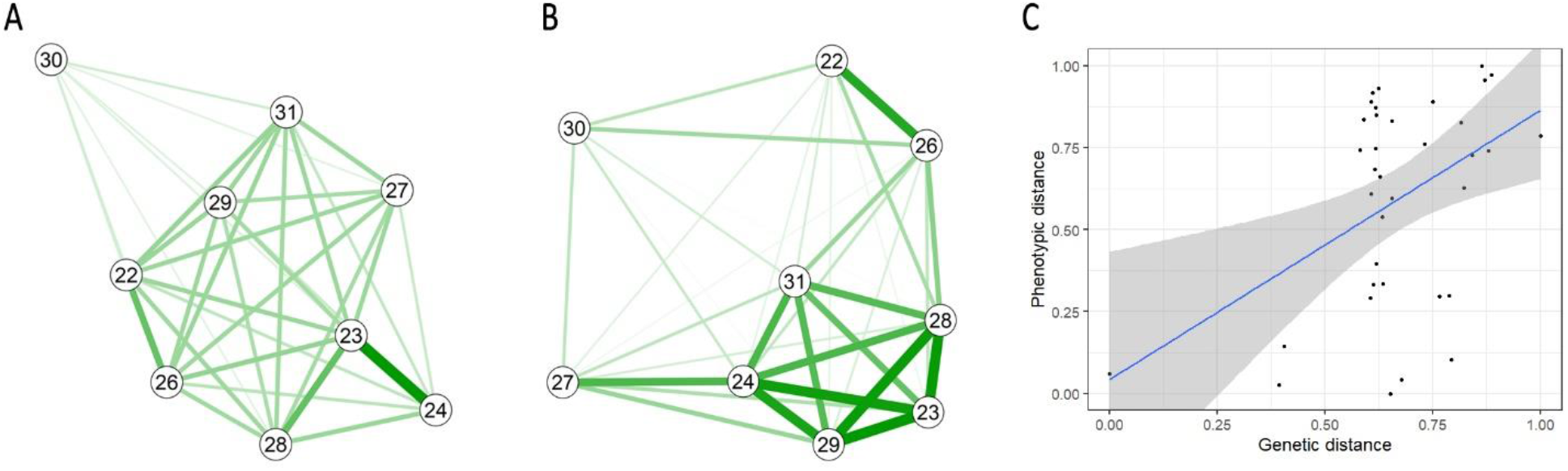
Phenotypic distances of nine average eggs laid by nine genotyped common cuckoo females (A) and their genetic distances (B). Thicker green lines denote higher phenotypic and genetic similarity. Correlation between phenotypic and genetic distances (C).

### Human assessment

Participants with some experience of working with biological data performed better at the grouping task than those with no experience, though there is no clear evidence that specific experience of working with cuckoo eggs is beneficial (Table 1).

**Table 1.** Cluster similarities of egg sorting performed by humans both without knowledge (when they did not know the number of females) and with a known number of females.

| Group | No knowledge | Known number of females |
| --- | --- | --- |
| No experience (n = 5) | 0.225 (0.066) | 0.241 (0.041) |
| Non-specific experience (n = 3) | 0.502 (0.170) | 0.496 (0.057) |
| Specific experience (n = 4) | 0.417 (0.050) | 0.456 (0.158) |

Mean cluster similarity (and SD in brackets) is presented for each category.

### Unsupervised learning

Clustering using unsupervised hierarchical learning gave a cluster similarity value of 0.452; similar to that of experienced human observers, but better than inexperienced observers.

### Supervised learning (random forest analysis)

#### Female clustering

Clustering using supervised random forest analysis (with a leave-one-out protocol) led to good classification, with a mean of 77.08/95 (81.1%) of eggs correctly assigned to their genetic parent. The cluster similarity had a mean of 0.61 (SD = 0.03), higher than both experienced human assessment and unsupervised learning.

We assessed variable importance (Table 2) using a full model including all data. PC2 for pattern was the most important variable for classification, and the variables loading onto this PC were predominantly those for the ‘skew’ of the pattern. PC2 for spectra was also important, with this variable being influenced by the ‘blueness/greenness’ of the egg. Finally, PC3 and PC1 for shape were also informative. The variables loading onto these PCs were the length, width, volume and surface area of the egg.

**Table 2:**
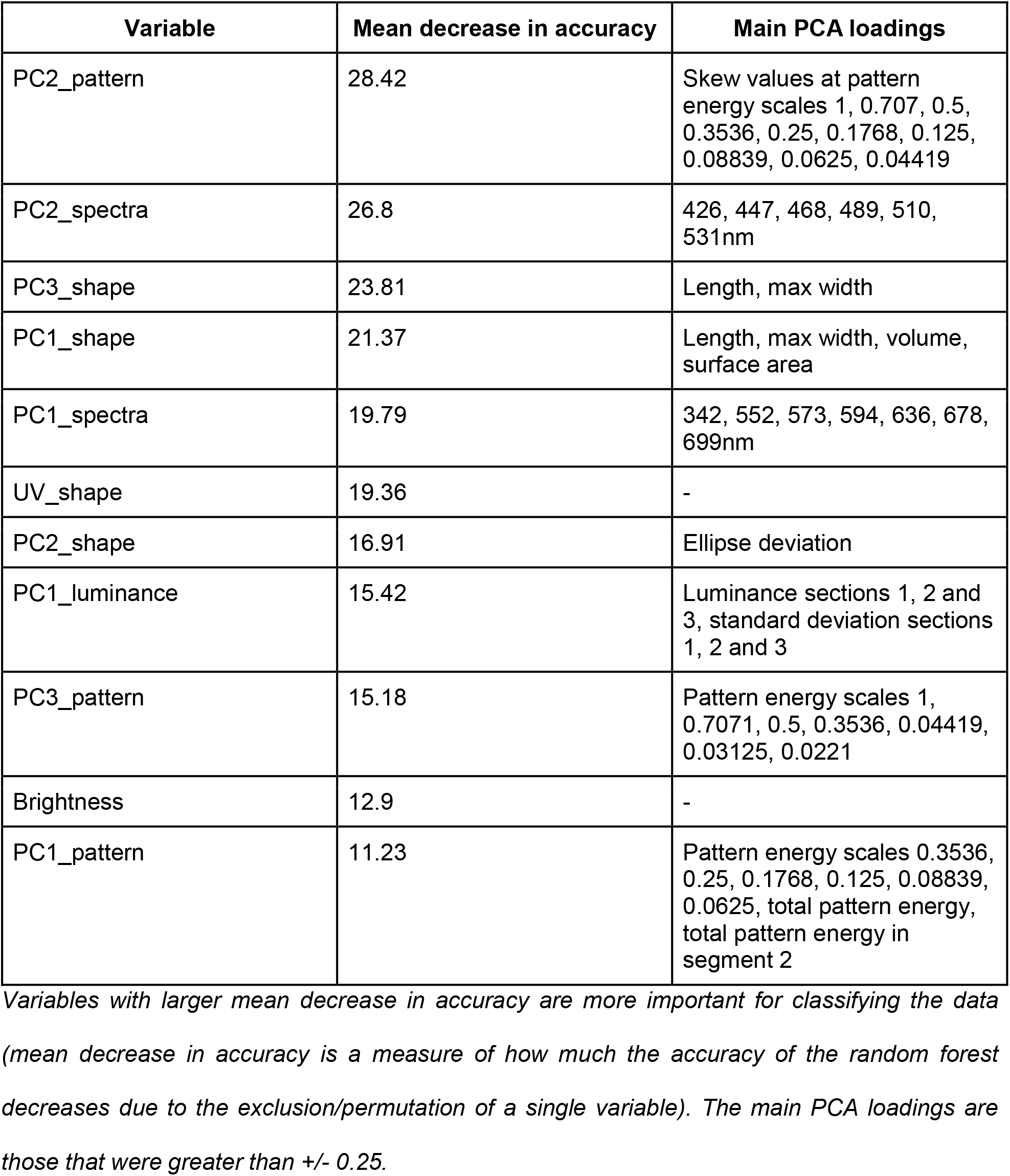
The importance of individual variables in egg clustering using random forest analysis.

| Variable | Mean decrease in accuracy | Main PCA loadings |
| --- | --- | --- |
| PC2_pattern | 28.42 | Skew values at pattern energy scales 1, 0.707, 0.5, 0.3536, 0.25, 0.1768, 0.125, 0.08839, 0.0625, 0.04419 |
| PC2_spectra | 26.8 | 426, 447, 468, 489, 510, 531nm |
| PC3_shape | 23.81 | Length, max width |
| PC1_shape | 21.37 | Length, max width, volume, surface area |
| PC1_spectra | 19.79 | 342, 552, 573, 594, 636, 678, 699nm |
| UV_shape | 19.36 | - |
| PC2_shape | 16.91 | Ellipse deviation |
| PC1_luminance | 15.42 | Luminance sections 1, 2 and 3, standard deviation sections 1, 2 and 3 |
| PC3_pattern | 15.18 | Pattern energy scales 1, 0.7071, 0.5, 0.3536, 0.04419, 0.03125, 0.0221 |
| Brightness | 12.9 | - |
| PC1_pattern | 11.23 | Pattern energy scales 0.3536, 0.25, 0.1768, 0.125, 0.08839, 0.0625, total pattern energy, total pattern energy in segment 2 |

*Variables with larger mean decrease in accuracy are more important for classifying the data* *(mean decrease in accuracy is a measure of how much the accuracy of the random forest* *decreases due to the exclusion/permutation of a single variable). The main PCA loadings are* *those that were greater than +/- 0.25.*

### Same/different analysis

For our labelled data, on average, 77.46/95 (81.5%) of eggs were matched with themselves. During the training phase, the model was always trained with ‘same’ pairs that consisted of two different eggs from the same female, so this was done as a test of whether the model was able to recognise two truly identical eggs as coming from the same female.

In addition, on average 73.50/95 (77.4%) eggs were uniquely matched with other eggs laid by the same female.

Out of 87 unlabelled eggs, the model was able to reliably (on 95% of runs) identify 22 as belonging to a labelled female (9 eggs assigned to female 5, 5 eggs to female 27, 3 eggs to female 13, and 1 egg to each of females 4, 21, 28, 29, 30).

### Phenotypic-genotypic similarity

There was no significant relationship between overall female egg phenotype distance and female genetic distance (Mantel test r = 0.1968, p = 0.098, Fig. 2).

When considering each aspect of phenotypic similarity separately, both pattern/luminance and shape distance metrics did not correlate with genotypic similarity (r = 0.0254, p = 0.3861 and r = −0.2317, p = 0.9256 respectively, Fig. 3A-D). However, spectral similarity did correlate with genetic similarity (r = 0.356, p = 0.037, Fig. 3E-F).

**Figure 3.**
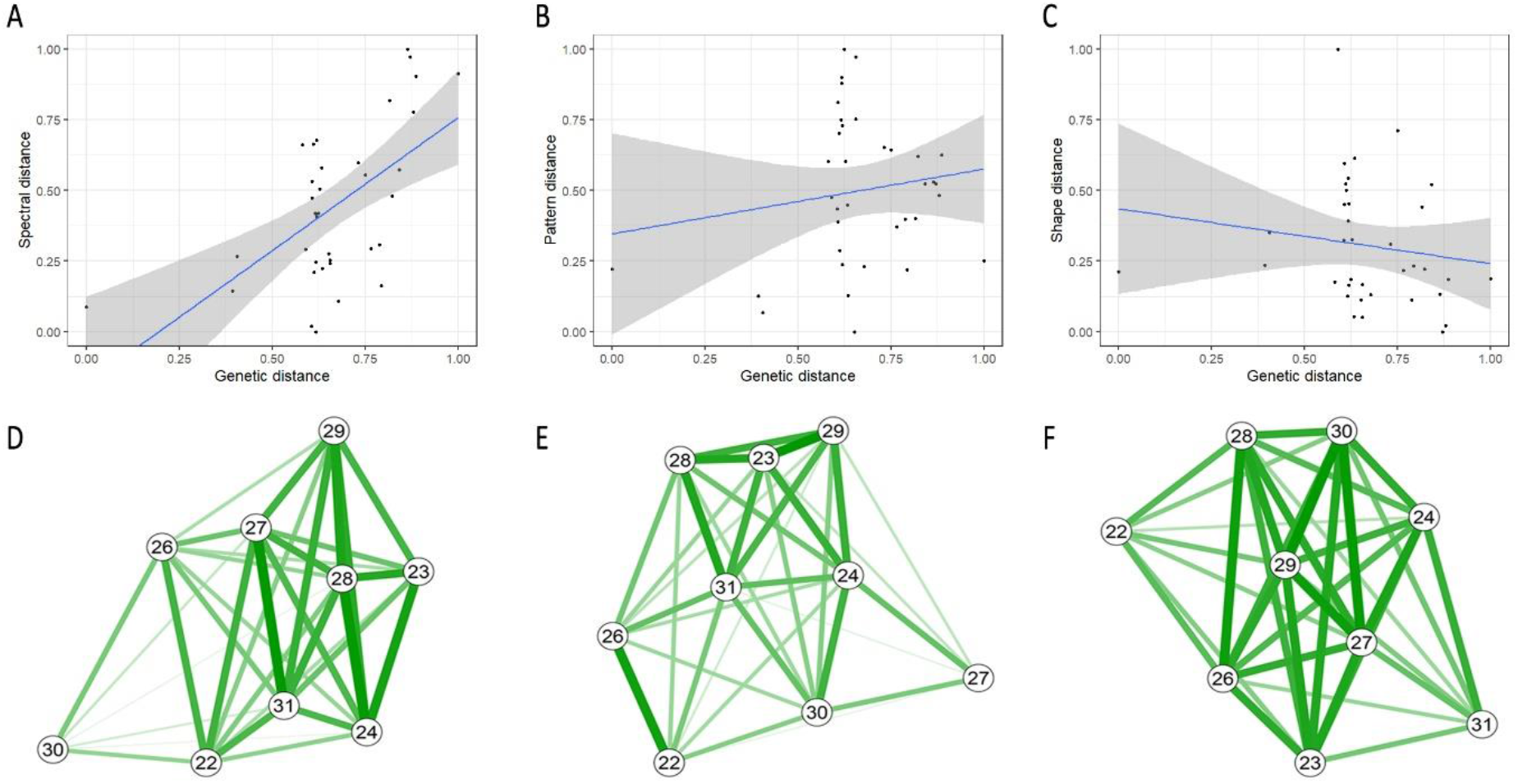
Correlation between spectral (A), pattern/lumiance (B) and shape (C) distances, respectively and genetic distances. Individual phenotypic distances of average eggs laid by nine genotyped common cuckoo females: spectral (D), pattern/luminance (E) and shape (F) distances.

## Discussion

The results of our study support the ‘constant egg-type hypothesis’ predicting that individual cuckoo females lay eggs with a relatively constant appearance (Latter 1902, Chance 1940, Baker 1942, Lack 1968, Wyllie 1981, Moksnes et al. 2008). We confirmed that the within-clutch variation of cuckoo egg appearance is significantly lower than between-clutch variation. This has also been observed in other bird species and several adaptive explanations have been proposed for this phenomenon, such as easier recognition of the parasitic egg by hosts (Øien et al., 1995), recognition of an individual’s own clutch in colonially-breeding birds (Hauber et al., 2019) or signalling the female quality (Moreno & Osorno, 2003). Moreover, it has been shown that diet has only a small effect on eggshell colour and that this trait is more affected by maternal identity, suggesting that egg colour may carry information about intrinsic properties of the female (Dearborn, Hanley, Ballantine, Cullum, & Reeder, 2012). Therefore, it seems that there is the potential to use egg appearance to identify individual bird females.

Here, we demonstrated that the unsupervised computer-vision based classifier can outperform human participants (especially inexperienced ones) in assigning cuckoo eggs to correct female clusters. Human egg classifiers with experience of handling natural eggs (either cuckoo or of other species) were able to more accurately sort them compared to people without this experience (Table 1). This may indicate that experienced observers have some knowledge of which cues are likely to be useful. They may also be more likely to be practiced at this type of fine discrimination task, or be more motivated to take part, given their interest in biology. Interestingly, knowing the number of groups (here: cuckoo females that laid eggs) did not increase sorting accuracy substantially.

The automatic hierarchical clustering method showed very similar results to experienced human classifiers, while supervised random forest analysis showed considerably better results and 81% of cuckoo eggs were assigned correctly. This suggests that in some cases, automatic assessment methods may be preferable to human assessment. Detailed consideration of the clusters created by humans and the automatic methods showed that the same females were problematic for both clustering methods, probably reflecting phenotypic overlap between some individuals (all sorting results can be found in Supplementary Material). Our results showed that one of the pattern characteristics (skew), blueness of colour and finally egg size were the most important parameters that helped the automatic method to cluster eggs the most accurately. The slight improvement in clustering accuracy for the automatic methods over human assessment may reflect the use of features that humans are not able to see (e.g. the UV peak of the spectra). Therefore, we would recommend automatic assessment over human assessment where possible.

A previous study suggested that closely related cuckoo females may lay eggs that are indistinguishable from each other (Moksnes et al., 2008). Our limited dataset of nine caught females for which we calculated genetic distances partially supports this idea as we found that the background colour of eggs was more similar between closely related females. However, pattern and dimension distances between females did not correlate with their genetic distances. As an interesting example, we caught two closely related females, mother (female 23) and daughter (female 24), and we photographed and measured the colour of their eggs. From the photographs, we can see that their eggs look very similar in the colour and spotting pattern; however, they differ considerably in size (Fig 3 in Supplementary material). Similarly, we can also see the resemblance in egg colour and pattern (but not in size) between other closely related females (females 23 vs 28 and females 22 vs 26; Fig. 3 and 4 in Supplementary material). Previous studies showed that all investigated egg features - colour, spotting pattern and also size have high heritability (Wei et al., 1992; Collias, 1993; Gosler et al., 2000; Christians, 2002; Morales et al., 2010). Our results indicate that the background colour might be a more heritable trait than spotting pattern and egg size or shape, which supports the idea that egg colour is of vital importance for egg recognition (Avilés et al., 2010; Spottiswoode & Stevens, 2010; Michal Šulc, Procházka, Capek, & Honza, 2016). Since several studies reported that hosts and even parasites themselves also use spotting pattern (López-de-Hierro & Moreno-Rueda, 2010; Spottiswoode & Stevens, 2010; de la Colina, Pompilio, Hauber, Reboreda, & Mahler, 2012) or egg size (Marchetti, 2000; Spottiswoode, 2013) when recognizing and eliminating parasitic eggs, we still expect relatively high heritability of these egg traits in brood parasites. We suspect that the insignificant relationship between genetic distance and phenotypic distance in spotting pattern and size is only a matter of our limited sample size. A Larger sample size, including more mother-daughter pairs, is needed to truly estimate heritability values of individual egg traits (de Villemereuil, Gimenez, & Doligez, 2013). The lack of significant correlation between egg shape and genetic similarity may also be explained by the fact that egg size often reflects the size of laying females (Larsson & Forslund, 1992; Nager & Zandt, 1994), which depends on the genetic contribution of both parents and therefore might differ more even in closely related females. Moreover, even within one host population, cuckoos are raised by host parents that vary in their provisioning care (Požgayová, Piálková, Honza, & Procházka, 2018), which may also influence the body size of cuckoo females in adulthood. Finally, some studies showed there is a positive relationship between food availability and egg size (reviewed in Christians 2002). Consequently, since egg size and shape may even differ in such closely related females, these traits may be used for the separation of their eggs. And indeed, this is what we observed in several of our human participants and also in computer analysis where separation of mother and daughter eggs was relatively straightforward presumably because of the apparent difference in the size and shape (see Supplementary material).

Although our results are in concordance with a previous study showing that the visual appearance of cuckoo eggs cannot be used to assign them reliably to individual females without genetic data (Moksnes et al. 2008), here we present a new approach that uses both genetic and phenotypic information. We used this method for assigning cuckoo eggs for which we did not have genetic data (because they were ejected by hosts or predated), allowing us to assign 22 eggs (out of 87) to eight known females. This method seemed to work well for females that laid very distinctive eggs and therefore results will strongly depend on within- and between-clutch variation. We may expect better results of the method in species where between-clutch variation substantially exceeds the within-clutch variation. It must also be noted that the accuracy of the assignment will increase with the relative number of (genetically and phenotypically) analysed samples in the study area that are used for the training dataset, because broad sampling will reduce the chance that an unsampled egg that has been laid by a completely new female will be assigned to an existing (incorrect) female. Finally, it is important to apply other information (laying date and laying area) to eliminate potential incorrect assignments. However, in our dataset, we did not find any such discrepancies for the 22 eggs that were automatically assigned based on their phenotypes.

We conclude that egg appearance alone cannot be used to identify individual cuckoo females. Clusters created either by people using egg photographs or by the computer using spectral and image data did not fully correspond to the true female identity acquired from molecular analyses, though the supervised automatic assessment was the most accurate for classification. Our results support the idea that more closely related females lay eggs more similar in their colour. However, the size and shape of the eggs seems to be the least heritable trait, which may substantially help to distinguish even between eggs of closely related cuckoo females. We advocate genetic analysis to be used for determination of maternity in this species. However, in sufficiently sampled systems, supervised analytical methods that use egg visual features might additionally help to broaden sample sizes, which may be very desirable for studying biological questions (see e.g. Koleček et al. 2020 in prep). We encourage researchers investigating inter- and intra-specific brood parasitism to use this low-cost and ethically more appropriate method of individual identification. Since a similar technique has been successfully used in non-parasitic species (Höltje et al., 2016), identification of laying females using egg appearance therefore has the potential to be of widespread use.

## Supporting information

Supplement_egg_photos

Supplement_ID_results

## Acknowledgements

We would like to thank Lisandrina Mari, Kamila Bendová, Kristýna Míčková, Jana Hodanová, Klára Stehlíková, Martina Fridrichová, Marta Potůčková, David Tesař, Václav Jelínek and Petr Potůček for their sorting of cuckoo eggs. Lisandrina Mari also substantially helped with writing of the genetic part of methods. We also thank Lukáš Kulísek, Boris Prudík, Kateřina Sosnovcová, Jaroslav Koleček and Miroslav Čapek for help with fieldwork. We thank Micha Elsner for his help with data analyses. We are also very grateful to Vladimír Beneš and the European Molecular Biology Laboratory Genomic Core Facility in Heidelberg (Germany) for their kind advice and technical support regarding Illumina sequencing. The managers of the Hodonín Fish Farm kindly permitted us conducting the fieldwork on their grounds.

## Ethical note

This study was carried out with the permission of the regional nature conservation authorities (permit numbers JMK: 115874/2013 and 38506/2016; MUHOCJ: 41433/2012/OŽP, 34437/2014/OŽP, and 14306/2016/OŽP). The fieldwork adhered to the animal care protocol (experimental project numbers 039/2011 AV ČR and 3030/ENV/17-169/630/17) and to the Czech Law on the Protection of Animals against Mistreatment (license numbers CZ 01272 and CZ 01284). This study was carried out with the permission of the regional nature conservation authorities (permit numbers JMK: 115874/2013 and 38506/2016; MUHOCJ: 41433/2012/OŽP, 34437/2014/OŽP, and 14306/2016/OŽP).

## Funding

This work was supported by the Czech Science Foundation (grant number 17-12262S) and by the Programme for research and mobility of young researchers of the Czech Academy of Sciences (MSM200931801, awarded to M.Š.).

## Notes

### Competing Interest Statement

The authors have declared no competing interest.

