## Supplement_egg_photos for "Automatic identification of bird females using egg phenotype"

**Supplementary material**

*Figures 1-5 Eggs assigned to individual cuckoo females based on genetic results (eggs without underscore) or based on phenotypic similarities using same-different analysis (red underscore). In brackets, RW denotes the reed warbler and GRW the great reed warbler host.*

*Figure 6 and 7 Eggs that were not genetically sampled and remained also unassigned by after same-different analysis.*

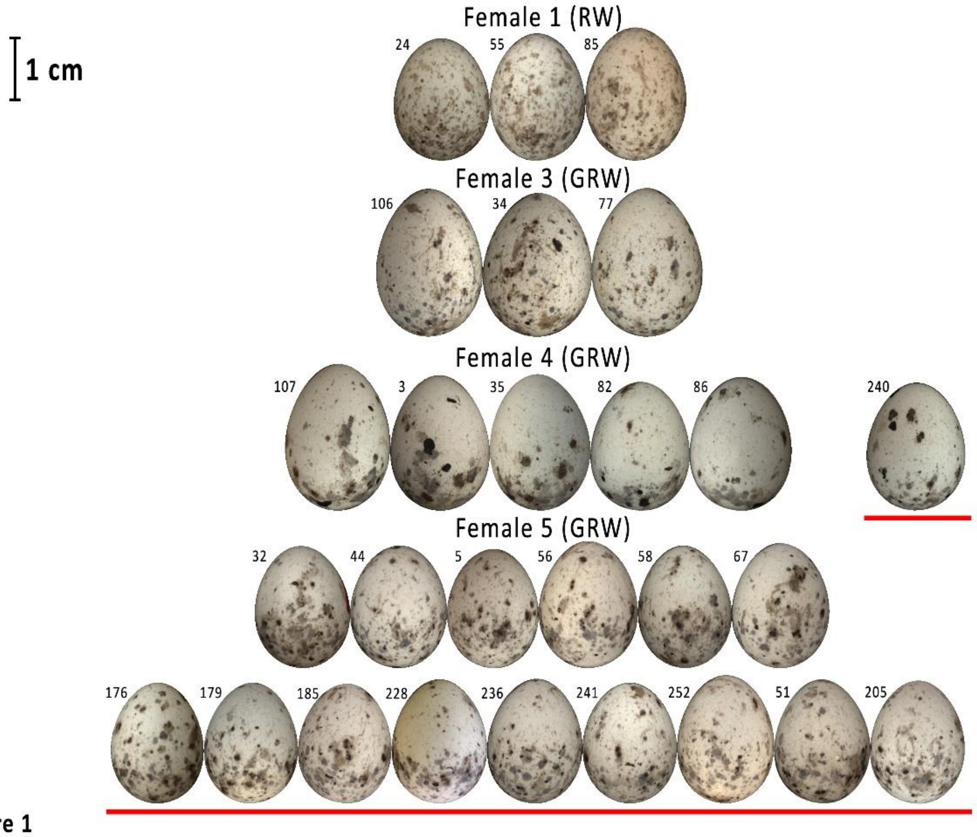


Figure 1

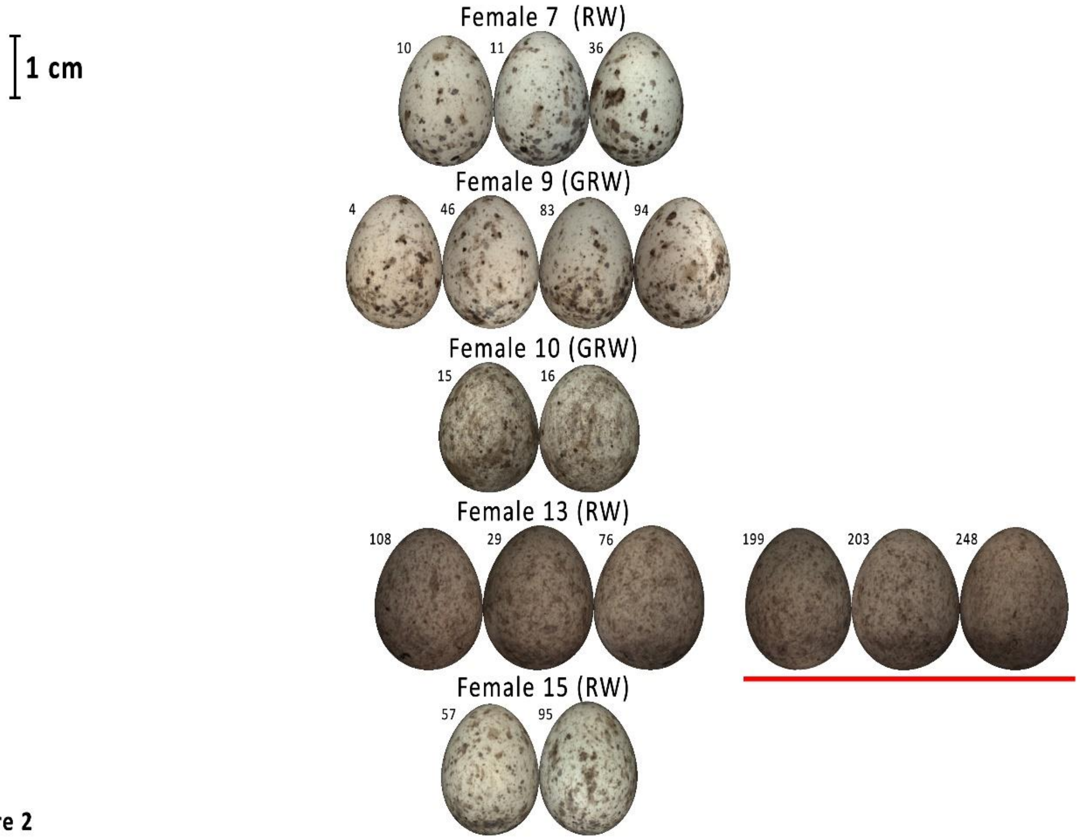


Figure 2

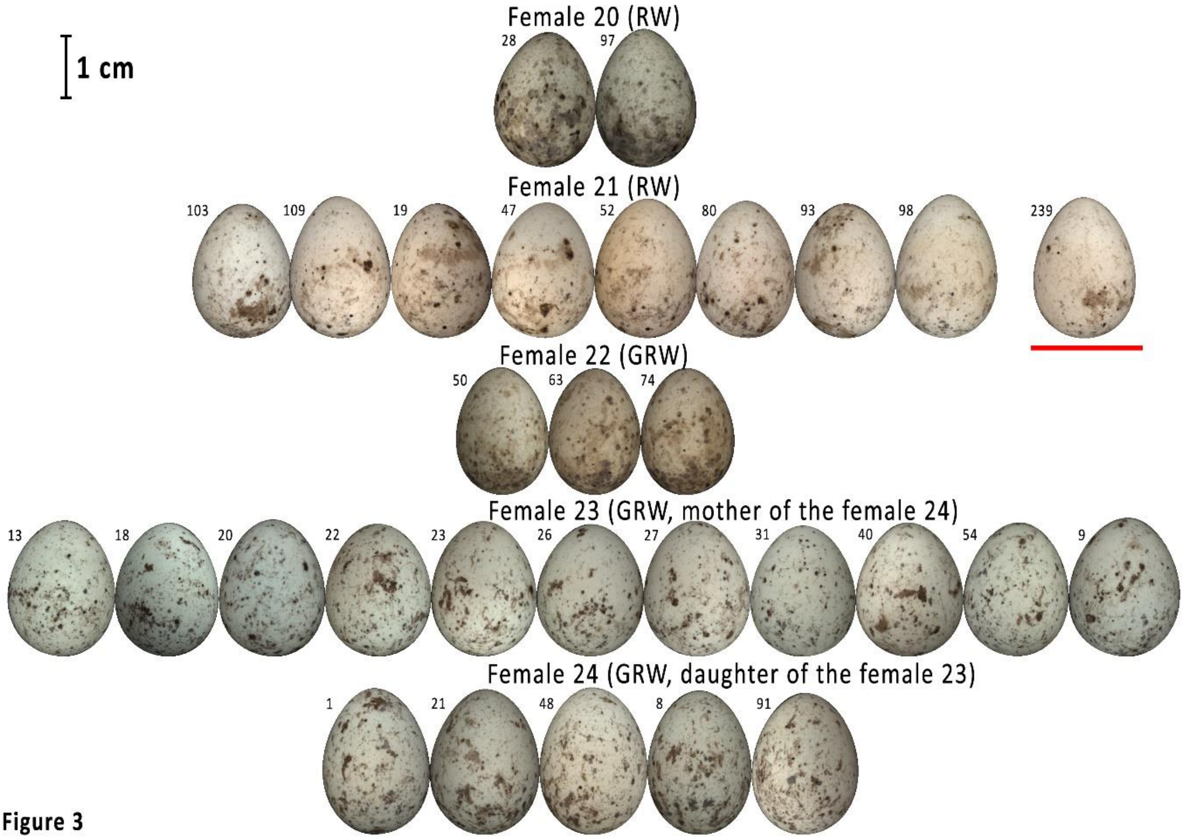


Figure 3

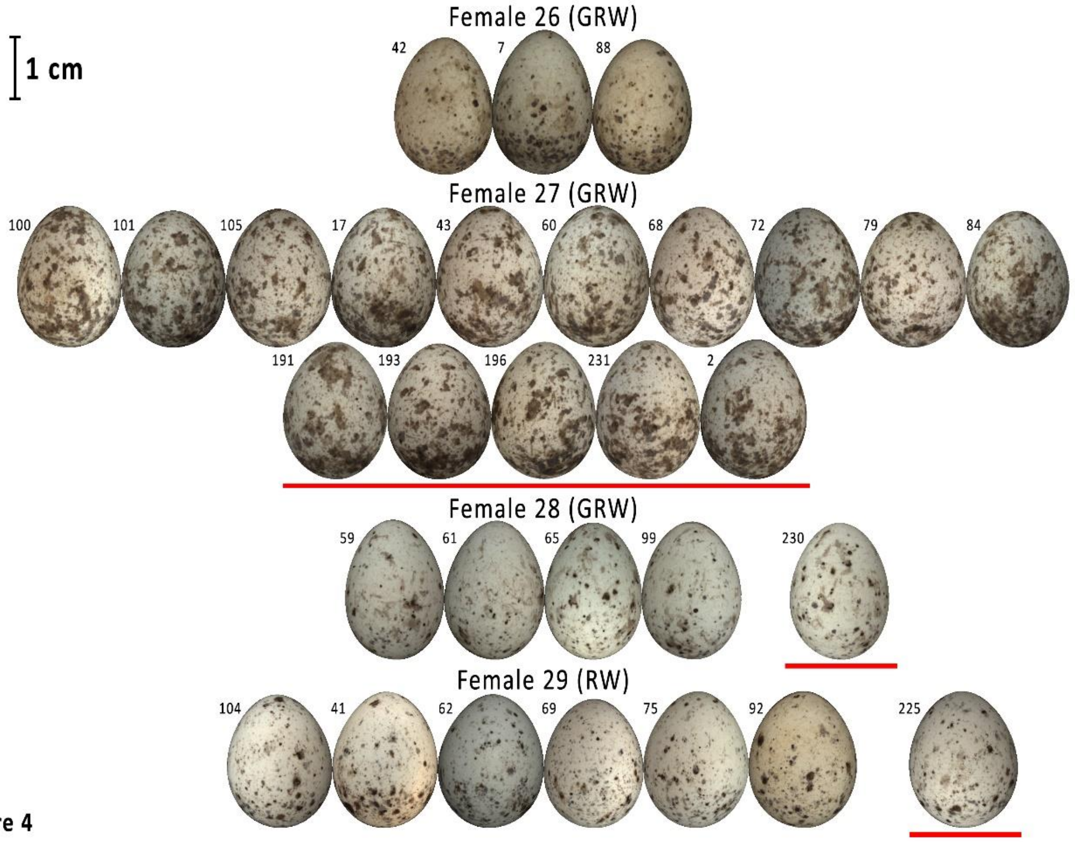


**Figure 4**

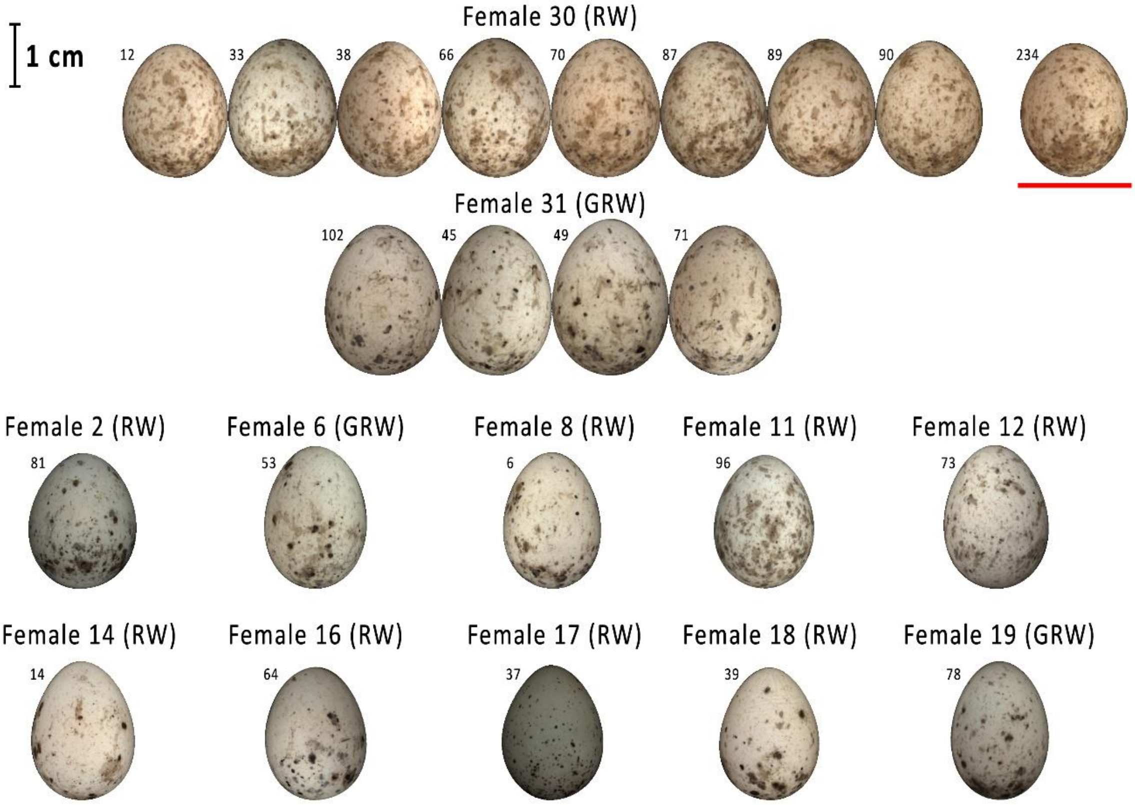


**Figure 5**

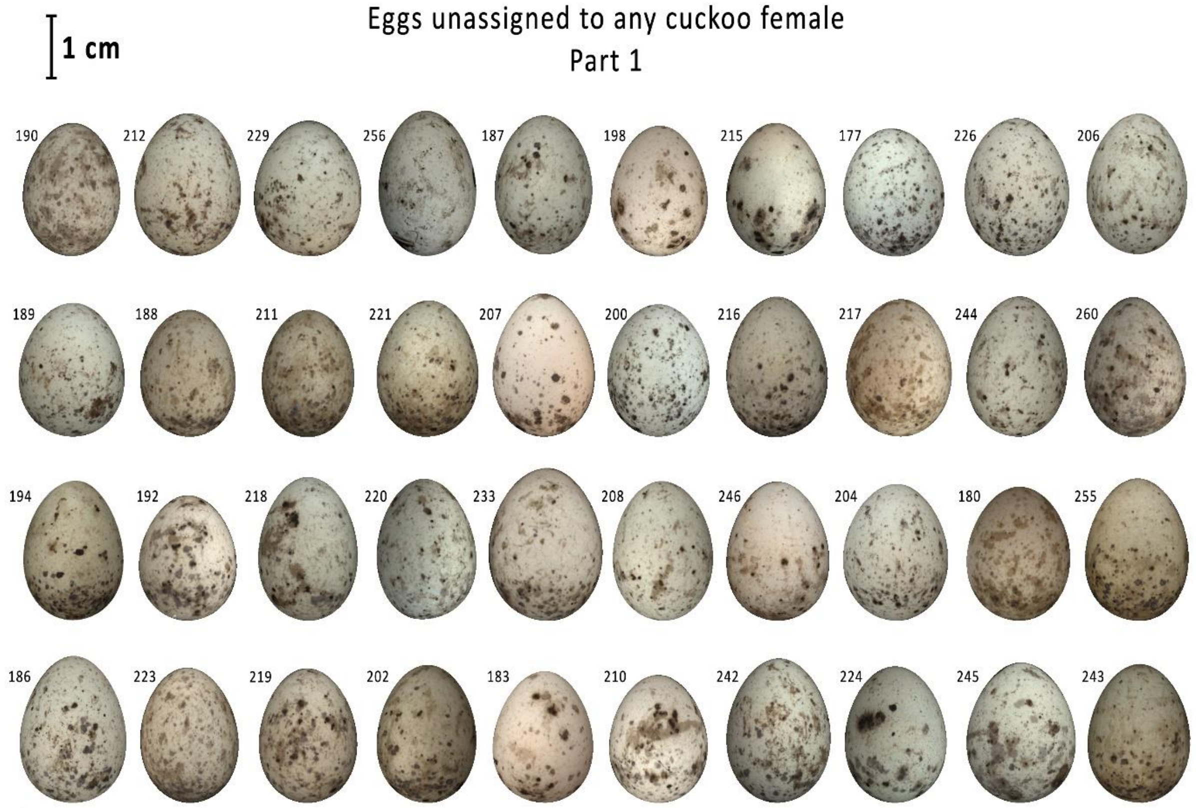


**Figure 6**

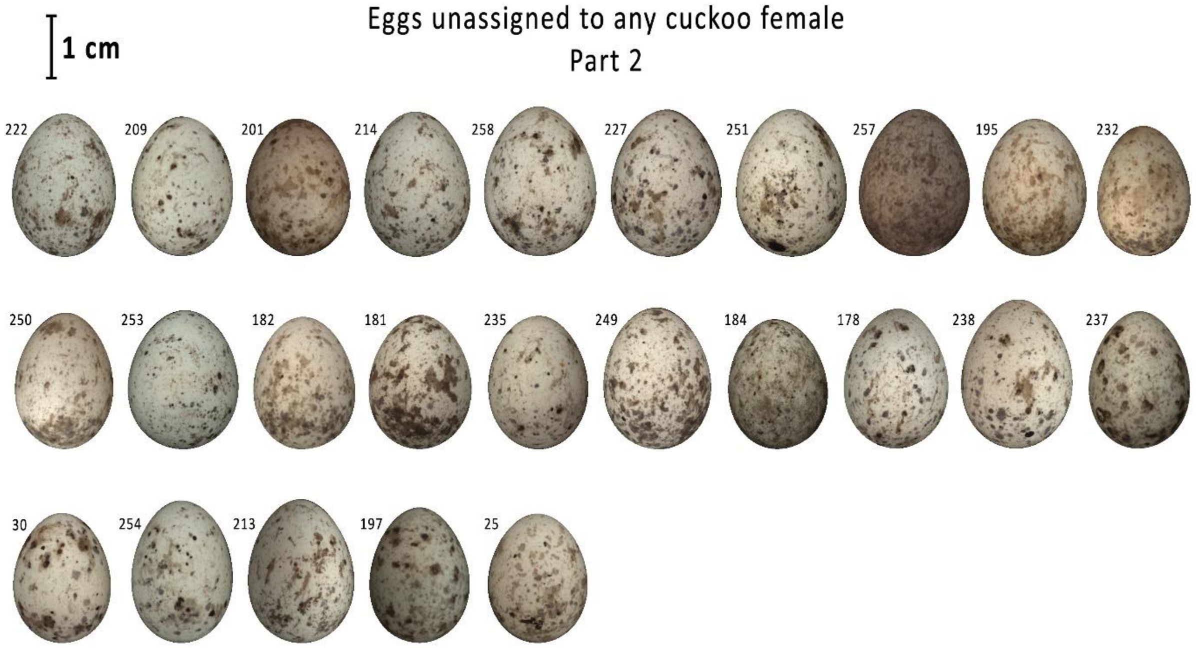


**Figure 7**
